# Conserved anti-inflammatory effects and sensing of butyrate in zebrafish

**DOI:** 10.1101/2020.05.13.069997

**Authors:** Pradeep Manuneedhi Cholan, Alvin Han, Brad R Woodie, Angela RM Kurz, Warwick J Britton, Lihua Ye, Zachary C Holmes, Jessica R McCann, Lawrence A David, John F Rawls, Stefan H Oehlers

## Abstract

Short chain fatty acids (SCFAs) are produced by microbial fermentation of dietary fiber in the gut. Butyrate is a particularly important SCFA with anti-inflammatory properties and is generally present at lower levels in inflammatory diseases associated with gut microbiota dysbiosis in mammals. We aimed to determine if SCFAs are produced by the zebrafish microbiome and if SCFAs exert conserved effects on zebrafish immunity as an example of the non-mammalian vertebrate immune system. We demonstrate that bacterial communities from adult zebrafish intestines synthesize all three main SCFA *in vitro*, although SCFA were below our detectable limits in zebrafish intestines *in vivo*. Immersion in butyrate, but not acetate or propionate, reduced the recruitment of neutrophils and M1-type pro-inflammatory macrophages to wounds. We found conservation of butyrate sensing by neutrophils via orthologs of the *hydroxycarboxylic acid receptor 1* (*hcar1*) gene. Neutrophils from Hcar1-depeleted embryos were no longer responsive to the anti-inflammatory effects of butyrate, while macrophage sensitivity to butyrate was independent of Hcar1. Our data demonstrate conservation of anti-inflammatory butyrate effects and identify the presence of a conserved molecular receptor in fish.

## Introduction

Short chain fatty acids (SCFAs) are microbial metabolites produced in the gut by the anaerobic fermentation of dietary fiber and protein in the large intestine ^1^. The most abundant SCFAs are acetate, butyrate, and propionate. In addition to providing the host with an energy source, microbially-derived SCFAs exert anti-inflammatory effects through inhibition of histone deacetylates (HDAC) and activation of G-protein coupled receptors (GPCRs) ^2^. Most research on SCFAs has reported their anti-inflammatory properties in mammals ^3-5^. However, the anti-inflammatory mechanism responsible for the anti-inflammatory effects of SCFA administration has not been reported in fish species to date.

Zebrafish are an important model of vertebrate gut physiology with key experimental advantages including high fecundity, transparency, and well-developed gut digestive function by 6 days post fertilization (dpf) ^6^. There is a high degree of intestinal immune conservation across vertebrates, including the sensitivity of zebrafish intestinal epithelial cell progenitors to butyrate ^7-9^. However, SCFA production has not been previously observed in the intestines of zebrafish, and it is unclear if the intestinal lumen of the zebrafish intestine provides a suitable niche for SCFA production ^9^. Nevertheless, detectable amounts of acetate, butyrate, and propionate have been measured in several species of teleosts ^10-14^.

We have used conservation between mammalian intestinal function and immunity to create zebrafish models of human intestinal inflammation ^15,16^. Key pattern recognition molecule families, such as the Toll-like receptors and Nod-like receptors, are evolutionarily ancient, and have conserved roles in zebrafish intestinal immunity ^7,8,17,18^. However, conservation of host molecules responsible for sensing SCFAs has not been explored in teleosts. Mammals utilize a wide range of molecules to sense SCFAs including G protein-coupled receptors (GPRs) GPR81 (also known as HCAR1) which is primarily present on immune cells and GPR109A which is expressed on intestinal epithelial cells. Microbially-derived SCFAs also exert direct effects on host physiology through histone deacetylase (HDAC) inhibition ^19-21^.

In this study, we investigated whether SCFAs are produced in the zebrafish intestine, and if the anti-inflammatory effects and sensing of SCFAs are conserved in zebrafish. We find that the pattern of SCFA production by the zebrafish intestinal microbiota is different from that seen in mammals, but that the anti-inflammatory effects mechanisms of butyrate are conserved across vertebrate species and development regardless of the ability of their endogenous microbiota to produce measurable butyrate.

## Methods

### 2.1 Zebrafish handling

Adult zebrafish were housed at the Centenary Institute (Sydney Local Health District AEC Approval 17-036) and Duke University School of Medicine ((Duke Institutional Animal Care and Use Committee Protocol Approval A115-16-05). Adult zebrafish experimentation was approved by the Institutional Animal Care and Use Committees of Duke University approval A115-16-05. Zebrafish adults were reared, housed, and fed as previously described ^22^. All zebrafish embryo research experiments and procedures were completed in accordance with Sydney Local Health District animal ethics guidelines under approval 17-036. Zebrafish embryos were obtained by natural spawning and embryos were maintained and raised in E3 media at 28°C.

### 2.2 SCFA quantification from adult zebrafish

Adult zebrafish were euthanized with 200 – 300 mg/L of ethyl 3-aminobenzoate methanesulfonate (tricaine) (Sigma, E10521) prior to dissection. For each sample, intestines dissected from five adult (90+ dpf) EK WT zebrafish males (roughly 0.2 g total) were pooled and homogenized using a Precellys 24 High-Powered Bead Homogenizer at 5500 rpm for 3 cycles at 20 seconds per cycle with a 10 second delay between cycles. Samples were then acidified with HCl to a pH below 3, pelleted by centrifugation, and the supernatant was harvested. Filtered supernatant was stored at -80°C until quantification.

SCFA quantification as carried out on an Agilent 7890B GC FID, with an HP-FFAP capillary column (25 m length, ID 0.2 mm, film thickness 0.33 µm). Concentrations were determined using a linear model fit of a standard curve that encompasses the sample concentration range. Standardized concentrations used for each C2-C5 SCFA were as follows: 0.2, 0.5, 1, 2, 4, and 8 mM.

### 2.3 In vitro synthesis of SCFA by zebrafish gut commensals

Freshly dissected intestines from four adult (6-month-old) EK WT zebrafish males and frozen mouse fecal pellets were homogenized under reducing conditions to preserve the anaerobes. Samples were handled in a Coy anaerobic chamber and used to inoculate tubes containing brain-heart infusion (BHI) media (Thermo Scientific, OXOID) or Gifu anaerobic media (Sigma), both supplemented with deoxygenated hemin and vitamin K to a final concentration of 12.5 mg/L of hemin and 2.5 mg/L of vitamin K.

Tubes were incubated in a sealed anaerobic jar with a Gas-Pak (Becton and Dickinson) to maintain anaerobic conditions at 28°C for 24 hours. Samples were then acidified to a pH below 3 with HCl, pelleted by centrifugation, and the supernatant was harvested. Filtered supernatant was stored at -80°C until quantification with methods identical to those listed above in Section 2.2.

### 2.4 Drug treatments

Embryos were treated with 30 mM sodium acetate (Sigma; S2889), 30 mM sodium butyrate (Sigma; B5887), 30 mM sodium propionate (Sigma; P1880), 50 µg/mL dexamethasone (Sigma; D4902), or 100 mM 6-aminocaproic acid (Sigma; A2504). Drug stocks were dissolved in DMSO or PBS and added to E3.

### 2.5 Tail wounding experiment

Caudal fin amputation was performed on 5 dpf embryos unless otherwise indicated. Zebrafish embryos were anesthetized with tricaine. Embryos were cut posterior to the notochord using a sterile scalpel. Embryos were then recovered to fresh E3 and kept at 28°C.

### 2.6 Imaging

Live zebrafish embryos were anesthetized using tricaine, mounted on 3% methylcellulose (Sigma, M0512), and imaged using a Leica M205FA. ImageJ software was used to quantify the fluorescent pixel count within 100 µm of the wound site.

Additional high resolution and time-lapse microscopy was carried out on anesthetized embryos embedded in 1% low melt agarose in a 96 well-plate with a Leica SP8 confocal microscope or Deltavision Elite microscope.

### 2.7 Neutrophil tracking

Time-lapse images were processed and analyzed using ImageJ. Neutrophils were tracked using the Trackmate plugin in ImageJ software and further quantified using Chemokine and Migration tool software (Ibidi).

### 2.8 Germ-free derivation and microdissection of embryos

Germ-free zebrafish were created and maintained as previously described ^23^. The gut and body of 5 dpf embryos were separated using a 25-gauge needle and added to Trizol LS (Invitrogen; 10296010) for RNA extraction.

### 2.9 RNA extraction, cDNA synthesis and quantitative PCR (qPCR)

10 - 20 zebrafish embryos were pooled and lysed using a 25-gauge needle in Trizol LS for RNA extraction. cDNA was synthesized using a High-capacity reverse transcription kit (ThermoFisher Scientific, 4368814). qPCR was carried out using Power UP SYBR green master mix (ThermoFisher Scientific, 4385610) on a CFX96 Real-Time system (BioRad). Primer pairs (5’-3’); *18s* TCGCTAGTTGGCATCGTTTATG and CGGAGGTTCGAAGACGATCA; *hcar1* CATCGTCATCTACTGCTCCAC and GCTAACACAAACCGCACA.

### 2.10 gRNA synthesis and CRISPR injections

gRNA templates for *hcar1-2* (5’-3’): Target 1 TAATACGACTCACTATAGGTACCGGCGGCTCGATTGGGTTTTAGAGCTAGAAATAGC, Target 2 TAATACGACTCACTATAGGAGCAACTCTCGCTTCACTGTTTTAGAGCTAGAAATAGC, Target 3 TAATACGACTCACTATAGGGATTCGAGAGATGTTACTGTTTTAGAGCTAGAAATAGC. gRNA was synthesized as previously described ^24^.

A 1:1 solution of gRNA and 500 µg/mL of Cas9 nuclease V3 (Integrated DNA Technology) was prepared with phenol red dye (Sigma, P0290). Freshly laid eggs were collected from breeding tanks and the solution was injected in the yolk sac of the egg before the emergence of the first cell with a FemtoJet 4i (Eppendorf).

### 2.11 Statistics

All statistical analyses (t-tests and ANOVA where appropriate) were performed using GraphPad Prism8. Outliers were removed using ROUT, with Q = 1%. All data are representative of at least 2 biological replicates.

## Results

### 3.1 *Zebrafish gut commensals are capable of producing SCFA* ex vivo

We initially tried to detect SCFAs in whole intestines and their contents dissected from conventionally-reared adult zebrafish using gas chromatography but levels of acetate, butyrate, or propionate were below our limit of detection of 0.00132 mmol SCFA per g of tissue.

We next sought to determine if the conventional zebrafish gut microbiota had the capacity to produce SCFAs using *ex vivo* culture on two rich medias, BHI and Gifu. We found that microbial communities cultured from conventionally-reared adult zebrafish intestines were able to synthesize acetate under both aerobic and anaerobic conditions (**Figure 1**). Butyrate and propionate were only detected under anaerobic conditions. The highest concentrations of SCFA were detected under anaerobic conditions in BHI where acetate, propionate, and butyrate were present in a roughly 90:5:5 ratio.

**Figure 1:**
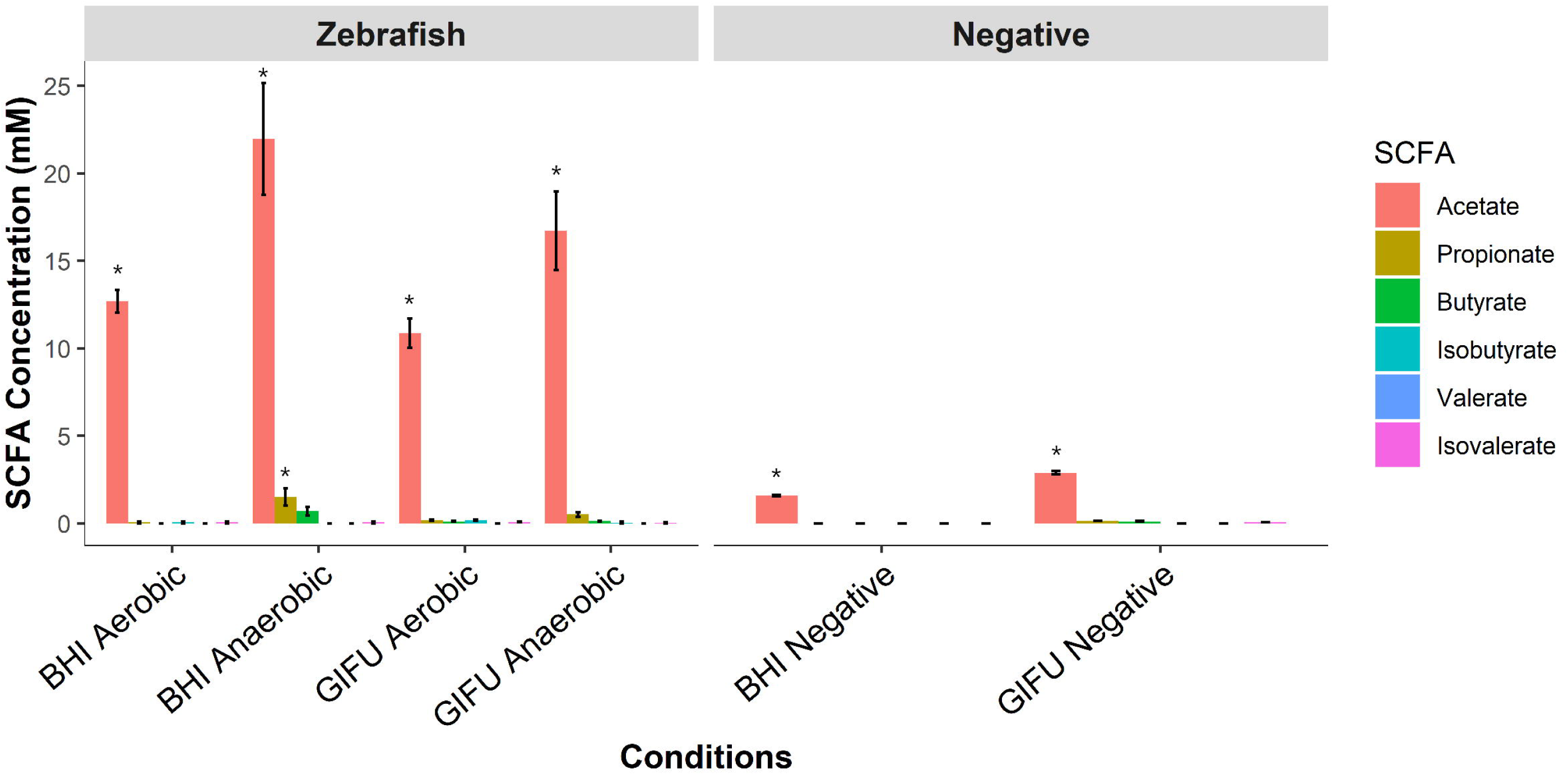
In vitro synthesis of SCFA by zebrafish gut microbiota. Concentrations of short-chain fatty acids (SCFA) synthesized by conventional microbiota harvested from adult zebrafish intestines. SCFA content of nutrient media used to culture microbes is provided under “Negative”. Asterisks indicate measurements within the range of standards. Error bars are shown as mean ± SE, n = 2. Bars without an asterisk indicate concentrations that were outside the standard range but were detectable.

### 3.2 Butyrate reduces the recruitment of zebrafish neutrophils to a wound

We first observed the effect of SCFAs (acetate, butyrate and propionate) on neutrophil migration following a tail wound injury using *Tg(lyzC:DsRed)*^*nz50*^ and *Tg(lyzC:GFP)*^*nz117*^ transgenic zebrafish lines where neutrophils are fluorescently labelled (**Figure 2A**). We observed a significant reduction in the number of recruited neutrophils at 6 hours post wounding (hpw) in embryos exposed to butyrate by immersion **(Figure 2B)**. There were no changes seen with acetate or propionate, but dexamethasone, a corticosteroid anti-inflammatory used as a positive control, reduced neutrophil recruitment as expected.

**Figure 2:**
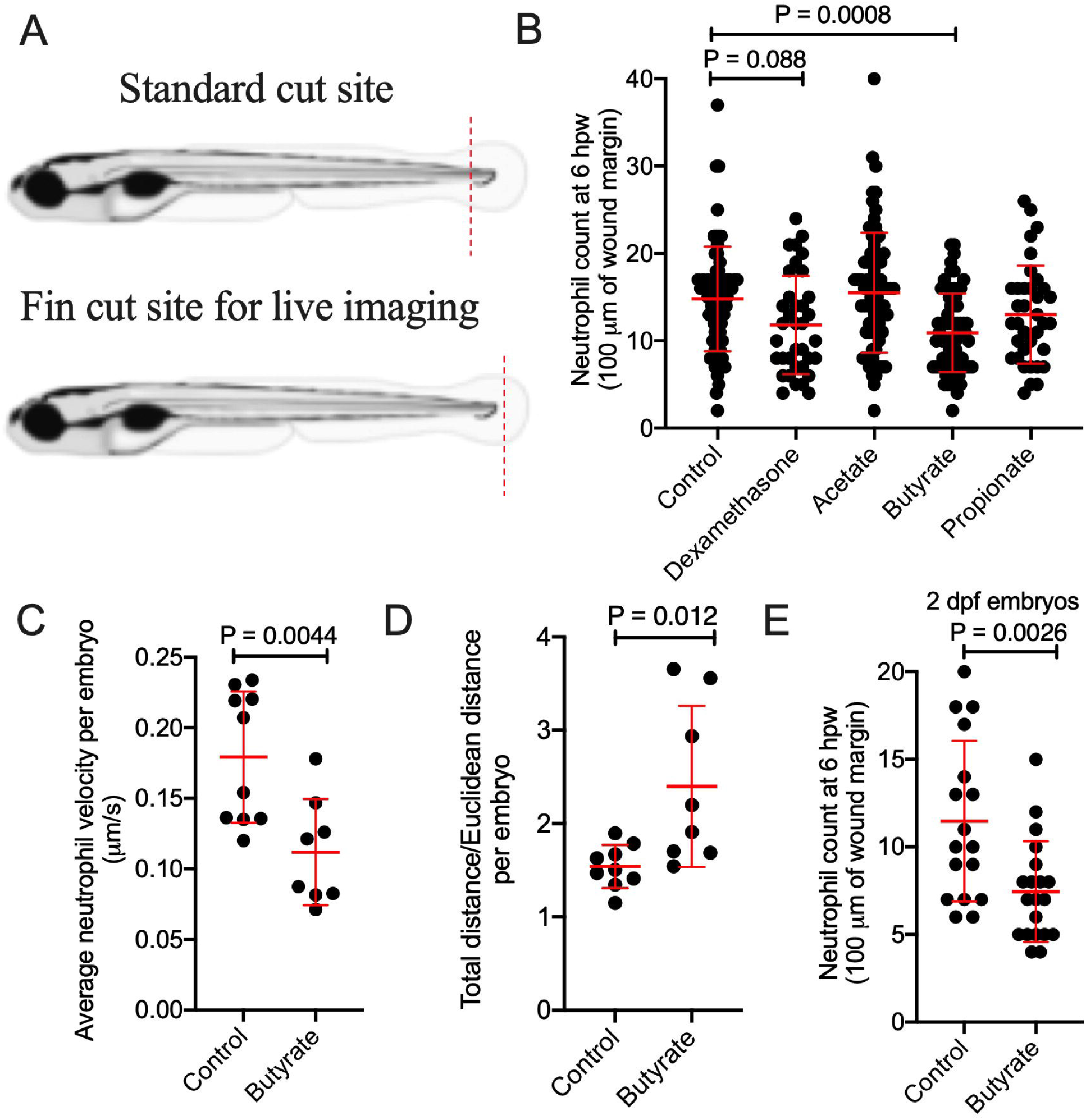
Butyrate reduces the recruitment of zebrafish neutrophils to a wound. (A) Cartoon describing the standard cut site transecting the dorsal aorta and cardinal vein of a 5 dpf zebrafish embryo, and the fin cut site used for live imaging studies. (B) Neutrophil counts at 6 hpw. Each dot represents a single embryo. (C) Velocity of wound-recruited neutrophils calculated from live imaging studies. Each dot represents the average of 10 neutrophils from a single embryo. (C) Meandering index (Total distance/Euclidean distance) of wound-recruited neutrophils calculated from live imaging studies. Each dot represents the average of 10 neutrophils from a single embryo. (D) Neutrophil count at 6 hpw in 2 dpf zebrafish.

We next assessed the quality of neutrophil recruitment by intravital imaging. We observed reduced neutrophil velocity (**Figure 2C**), and increased meandering index (total distance traveled / Euclidean distance) in butyrate-treated embryos (**Figure 2D**).

We next sought to determine if butyrate sensitivity is dependent on intestinal maturity by repeating the tail wound experiment using 2 dpf embryos, a developmental stage prior to significant intestinal morphogenesis ^25^, and found neutrophil recruitment was overall reduced compared to 5 dpf, but still further inhibited by butyrate immersion **(Figure 2E)**.

### 3.3 Butyrate reduces the proportion of TNF positive macrophages at the wound site

Next, we examined the effect of SCFAs on macrophage recruitment and polarization following wounding. *Tg(mfap4:tdTomato)*^*xt12*^ transgenic zebrafish were used to visualize macrophage numbers ^26^. Relative to control embryos, macrophage recruitment was reduced by butyrate and increased by propionate treatment at 6 hpw **(Figure 3A)**. Consistent with a lack of effect on neutrophil recruitment, acetate treatment did not affect the number of recruited macrophages, and as the positive control anti-inflammatory dexamethasone significantly reduced macrophage recruitment. These changes were maintained at 24 hpw when inflammation is in the resolution phase **(Figure 3B)**.

**Figure 3:**
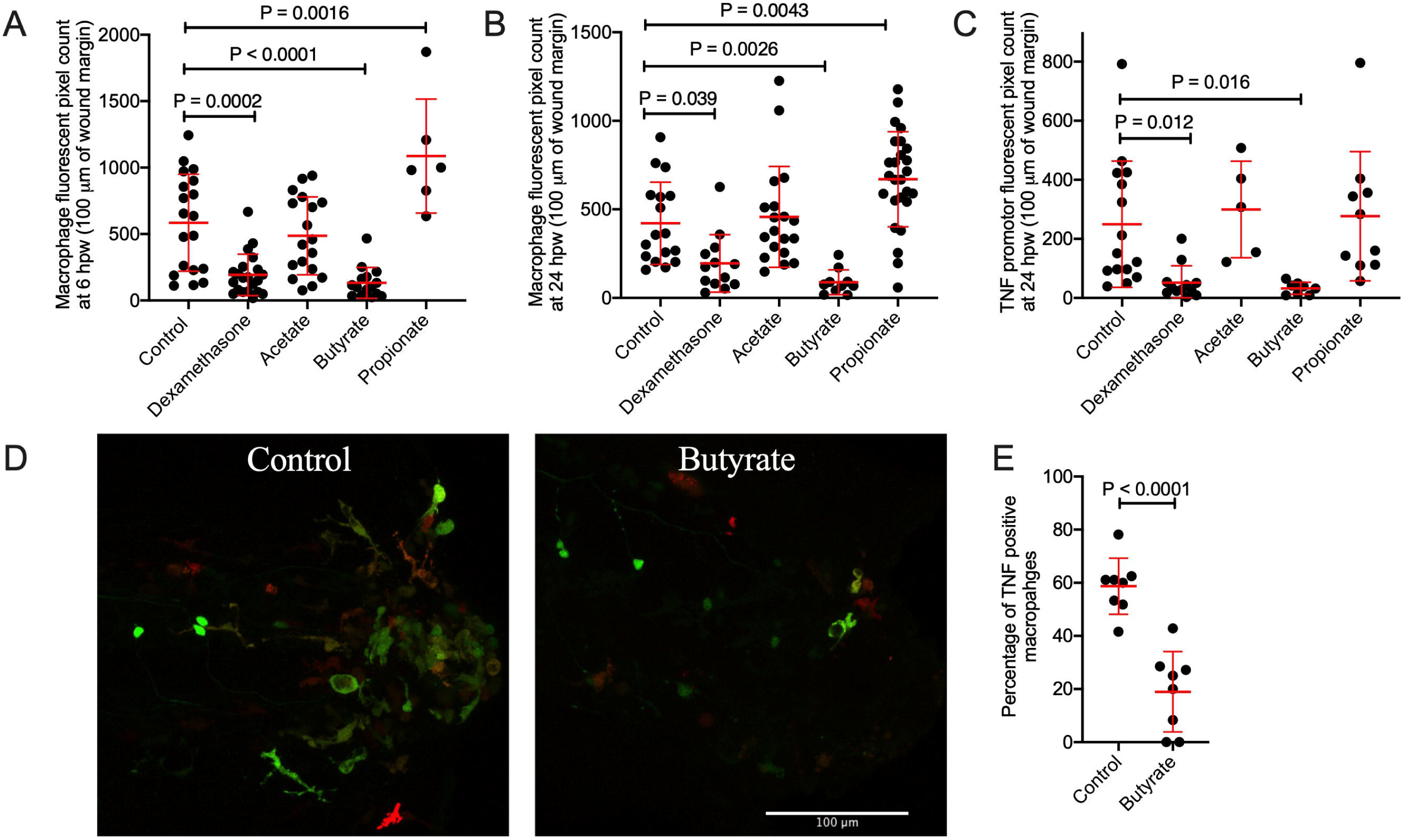
Butyrate reduces macrophage recruitment to the wound site and pro-inflammatory differentiation. (A) Macrophage fluorescent area at 6 hpw. (B) Macrophage fluorescent area at 24 hpw. (C) Total TNF promotor fluorescent area at the wound site after 24 hpw. (D) Representative images of double transgenic red macrophage, green TNF promoter activity embryos tail wounds at 24 hpw. Scale bar represents 100 µm. (D) Quantification of wound site TNF expressing macrophages at 24 hpw.

We next used the *TgBAC(tnfa:gfp)*^*pd1026*^ line to monitor inflammatory gene expression ^27^. As expected, since macrophages are the primary producers of TNF in zebrafish embryos, total TNF promotor fluorescence area at 24 hpw reflected the trend of macrophages recruitment in each treatment condition with butyrate, but not acetate or propionate, reducing TNF promoter expression **(Figure 3C)**. We next crossed the *TgBAC(tnfa:gfp)*^*pd1026*^ and *Tg(mfap4:tdTomato)*^*xt12*^ lines to monitor macrophage inflammatory polarization as defined by inflammatory TNF promotor expression (**Figure 3D**) ^28^. Butyrate treatment reduced the percentage of *TgBAC(tnfa:gfp)*^*pd1026*^ positive macrophages **(Figure 3E**).

### 3.4 Butyrate does not have toxic effects as measured by hemostatic indices

Butyrate has been previously shown to reduce the proliferation of zebrafish intestinal epithelial cells ^9^. This raises the possibility that inhibition of leukocyte recruitment by butyrate immersion was due to toxicity. Changes to zebrafish hemostasis have been observed in models of toxicity and inflammation ^29,30^.

We used *Tg(fabp10a:fgb-EGFP)*^*mi4001*^, where fibrin clots are visualized by GFP deposition, and *Tg(−6.0itga2b:eGFP)*^*la2*^, where thrombocytes are GFP-labeled, transgenic zebrafish lines to monitor hemostasis following transection of the dorsal aorta and posterior cardinal vein ^31,32^. We stabilized clots with aminocaproic acid as a positive control ^33^. Fibrinogen accumulation in *Tg(fabp10a:fgb-EGFP)*^*mi4001*^ embryos was unchanged at the wound site in response to butyrate treatment, however we noted that propionate treatment caused increased fibrinogen accumulation **(Supplementary Figure 1A)**. No changes were observed in thrombocytes accumulation in the *Tg(−6.0itga2b:eGFP)*^*la2*^ line following any of the SCFA treatments **(Supplementary Figure 1B)**.

### 3.5 Characterization of the zebrafish hydrocarboxylic acid receptor 1

*Hydrocarboxylic acid receptor 1* (*HCAR1*) is an important receptor for butyrate in mammals ^19^. We identified a conserved region of zebrafish chromosome 10, human chromosome 12, and mouse chromosome 5 containing *density regulated re-initiation and release factor (denr), coiled-coil domain-containing 62* (*ccdc62*), *huntingtin interacting protein 1 related a (hip1ra*) loci and the single exon *hcar* family (**Figure 4A**). Two copies of the putative zebrafish *hcar1* (with 93% identity and lacking sufficient divergence to differentiate by PCR) were identified as annotated by the single-entry NM_001163295.1 (Danio rerio hydroxycarboxylic acid receptor 1-2 (hcar1-2), mRNA) suggesting the possibility of a tandem duplication event (Supplementary File). Kuei *et al*. previously annotated what we refer as *hcar1a* as *gpr81-2* and *hcar1b* as *gpr81-1* ^34^. The predicted 933/936 bp transcript of *hcar1a/hcar1b* is expected to give rise to a 310/311 amino acid protein with 89% amino acid identity. The Hcar1a/Hcar1b hypothetical proteins contain 7 predicted transmembrane helix domains characteristic of a GPCR and an E-value of 7.05^-157^/3.21^-166^ for the HCAR subfamily ^35^. The predicted zebrafish proteins have approximately 43% identity to the human HCAR1, HCAR2, and HCAR3; and mouse HCAR1 and HCAR2 proteins.

**Figure 4:**
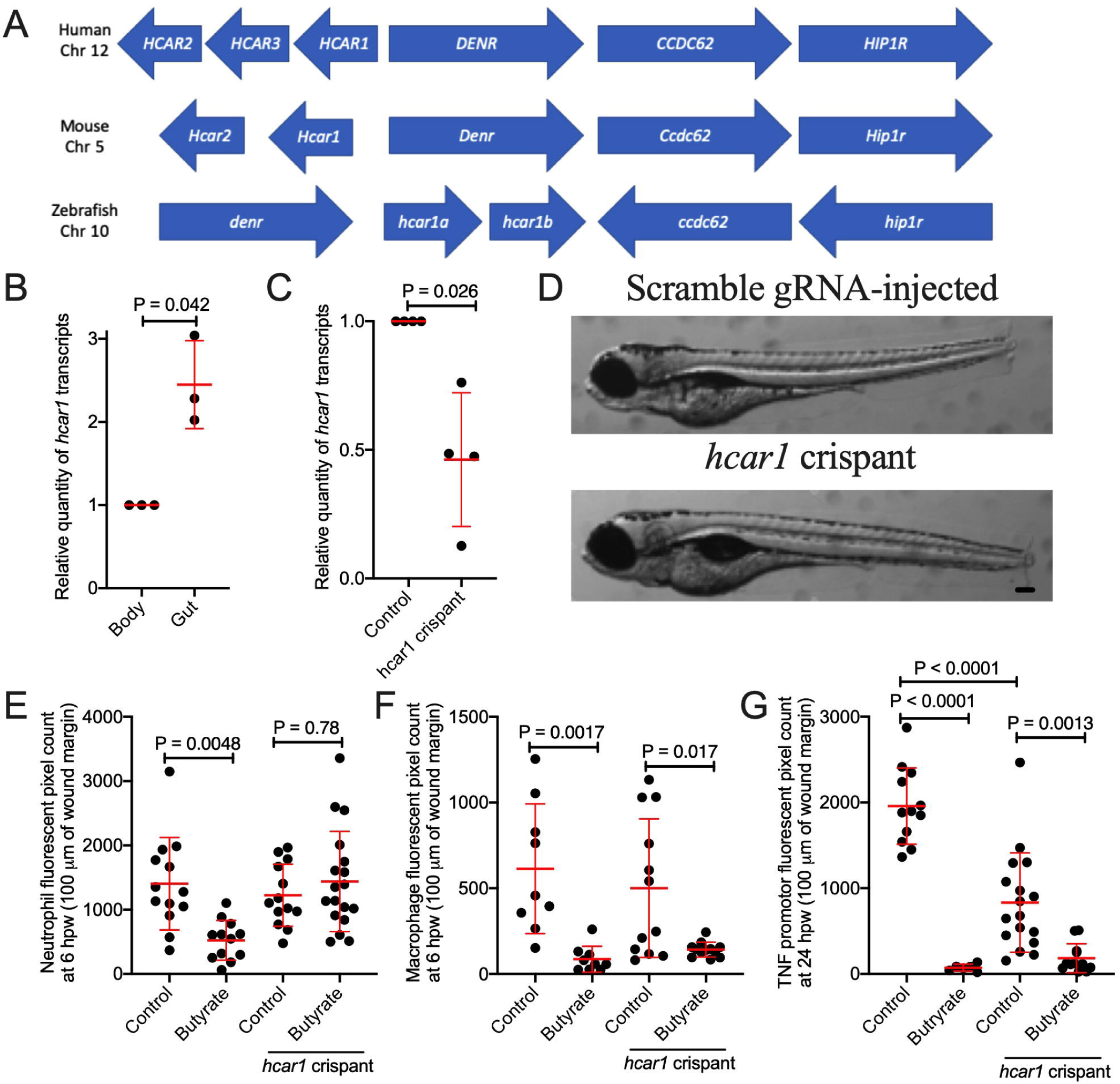
Characterization of zebrafish hydrocarboxylic acid receptor 1 and responsiveness to butyrate. (A) Synteny diagram illustrating *HCAR1* in a conserved region of human chromosome 12, mouse chromosome 5, and zebrafish chromosome 10. (B) Quantification of *hcar1* expression in dissected gut and body of zebrafish embryos. Each dot represents a biological replicate of at least 10 embryos. (C) Quantification of *hcar1* expression in 5 dpf embryos injected with *hcar1*-targeting Crispr-Cas9 complexes. Each dot represents a biological replicate of at least 10 embryos. (D) Morphology of the control and crispant embryos. Scale bar represents 100 µm. (E) Quantification of neutrophil area at 6 hpw in control and crispant embryos exposed to butyrate by immersion. (E) Quantification of macrophage area at 6 hpw in control and crispant embryos exposed to butyrate at 6 hpw. (F) Total TNF promotor-driven fluorescent area at the wound site at 24 hpw.

Mammalian *HCAR1* is highly expressed by intestinal epithelial cells and neutrophils ^19^. Compared to other immune cells, *hcar1* expression is enriched in zebrafish neutrophils ^36,37^. We sought to characterize expression of *hcar1* in the intestines of zebrafish embryos using microdissection of 5 dpf embryos. We found increased *hcar1* expression in dissected guts compared to the rest of the embryo by RT-qPCR analysis **(Figure 4B)**. Interestingly, there were no significant changes observed with the absence of microbial colonization in germ-free embryos (**Supplementary Figure 2**).

### 3.6 The anti-inflammatory effects of butyrate on neutrophils, but not macrophages, are dependent on Hcar1

To determine if butyrate acts through the Hcar1 receptor, we next used CRISPR/Cas9 technology to knockdown *hcar1* expression in zebrafish embryos. We utilized three target sites in *hcar1a*, two of which had strong homology to sequences in *hcar1b* (Supplementary File). We confirmed ∼50% transcript depletion by RT-qPCR for a shared *hcar1a/hcar1b* sequence (**Figure 4C**). Embryo development was morphologically normal compared to embryos injected with control scrambled guide RNA-Cas9 complex (**Figure 4D**).

We performed tail wound injury on *hcar1* knockdown embryos. Knockdown of *hcar1* abrogated the effect of butyrate treatment on neutrophil recruitment (**Figure 4E**). As expected from the lack of expression on macrophages, *hcar1* knockdown did not affect the reduced macrophage recruitment **(Figure 4F)** and TNF promotor expression **(Figure 4G)** induced by butyrate immersion.

## Discussion

This study shows for the first time that commensal microbiota residing in the zebrafish intestine are capable of producing SCFAs. Experimentally, we demonstrate that the effects, and sensing, of butyrate are conserved between zebrafish and mammals. Out of the three main SCFAs, only the anti-inflammatory effect of butyrate was found to be conserved in zebrafish embryos. We applied the commonly used tail wounding model to demonstrate an anti-inflammatory effect of butyrate on zebrafish neutrophils and macrophages. Using Crispr-Cas9 targeted mutagenesis, we also identified conserved butyrate responsiveness of the zebrafish Hcar1 receptor.

The anti-inflammatory effects of butyrate have been established in numerous *in vivo* and *in vitro* studies of mammalian hosts but not in fish ^38^. Our zebrafish tail wound model demonstrates conservation of this property in a bony fish. Bony fish diverge from mammals approximate 420 million years ago suggesting that the sensing of microbially-derived SCFAs has been conserved from a common ancestor.

Our finding that immune cells are responsive to butyrate even before intestinal lumen formation in early embryonic development is surprising as 2 dpf embryos are usually contained within relatively impervious chorions that prevent microbial colonization of the embryo. This suggests that the ability to sense xenobiotic SCFAs is programmed alongside the ability to sense more traditional microbially-associated molecular patterns via pattern recognition molecules.

The HCAR1/GPR81 butyrate receptor is expressed by many mammalian innate immune cells ^19^. Expression in zebrafish is strongest in granulocytes ^36,37^. Our knockdown experiments further demonstrate Hcar1 is necessary for the butyrate sensitivity of neutrophils but not macrophages. Thus, the butyrate-Hcar1-neutrophil behavior axis is evolutionarily ancient.

Human macrophages in the presence of butyrate have been shown to differentiate into an M2 phenotype which exhibits anti-microbial and tissue reparative properties ^39,40^. These effects are independent of HCAR1 signal transduction and are believed to be an effect of butyrate acting as a histone deacetylase inhibitor ^41,42^. SCFAs can permeate cell membranes through passive diffusion or through specific transporters such as the proton-coupled monocarboxylate-transporter 1 (MCT1) and sodium-coupled monocarboxylate-transporter 1 (SMCT1) ^43-46^. Consistent with this literature, we show butyrate reduces the expression of pro-inflammatory TNF by zebrafish macrophages independent of Hcar1 expression. Our data suggests the HCAR1-independent immunosuppressive actions of butyrate may be conserved across vertebrate evolution.

Our data demonstrate that, under nutrient-rich *in vitro* conditions, gut commensal microbiota from adult zebrafish are capable of synthesizing the three most important SCFAs: acetate, propionate, and butyrate. However, the ratio of acetate, propionate, and butyrate produced under anaerobic conditions in BHI media (90:5:5) differed from the ratio typically observed in mammalian colons (60:20:20) ^1^. This may be due to the differing bacterial communities found in zebrafish and mammalian intestines. The most abundant bacterial phyla in the adult zebrafish intestine are Proteobacteria and Fusobacteria, whereas mouse and human intestines are dominated by members of phyla Bacteroidetes and Firmicutes ^47^. Considering the SCFA production we observed *in vitro*, our inability to detect SCFA *in vivo* was surprising. We anticipate this could be due to rapid host or microbial metabolism of SCFA produced within the zebrafish gut, or the composition of the diet fed to the zebrafish tested in this experiment. Zebrafish are omnivores and were fed protein rich diets in the Duke aquaculture facility. A diet with more SCFA substrates such as carbohydrates and fiber may yield detectable SCFA production *in situ*.

Interestingly, we observed increased macrophage and fibrinogen clot accumulation at the wound site following propionate treatment, indicative of a pro-inflammatory effect. Although this is at odds with anti-inflammatory effects of propionate in mammals ^48-50^, it is consistent with evidence of an immunostimulatory effect of propionate in teleosts ^51,52^.

Overall this manuscript provides further evidence of conserved mechanisms of host-microbe interaction within vertebrates. We present evidence that immunological sensitivity to butyrate is conserved across vertebrates. Furthermore, there is conservation of the molecular machinery that senses butyrate even down to the responsiveness of individual leukocyte lineages.

## Funding

Australian National Health and Medical Research Council Project Grant APP1099912; The University of Sydney Fellowship G197581; NSW Ministry of Health under the NSW Health Early-Mid Career Fellowships Scheme H18/31086 to SHO.

The authors acknowledge the facilities and the scientific and technical assistance of the BioImaging Facility and Sydney Cytometry at Centenary Institute.

The authors declare no conflicts of interest.

## Figure Legends

**Supplementary Figure 1: Effects of SCFA administration on zebrafish hemostasis**.

(A) Clotting at 2 hpw following a tail wound in 5 dpf zebrafish. (B) Thrombosis at 3 hpw following a tail wound in 5 dpf zebrafish.

**Supplementary Figure 2: Expression of *hcar1* in germ-free embryos**

Quantification of *hcar1* expression in guts and bodies dissected from conventionally raised and germ-free embryos.

