## Supplementary figures and images for "Conserved anti-inflammatory effects and sensing of butyrate in zebrafish"

### Supplementary Figure 1

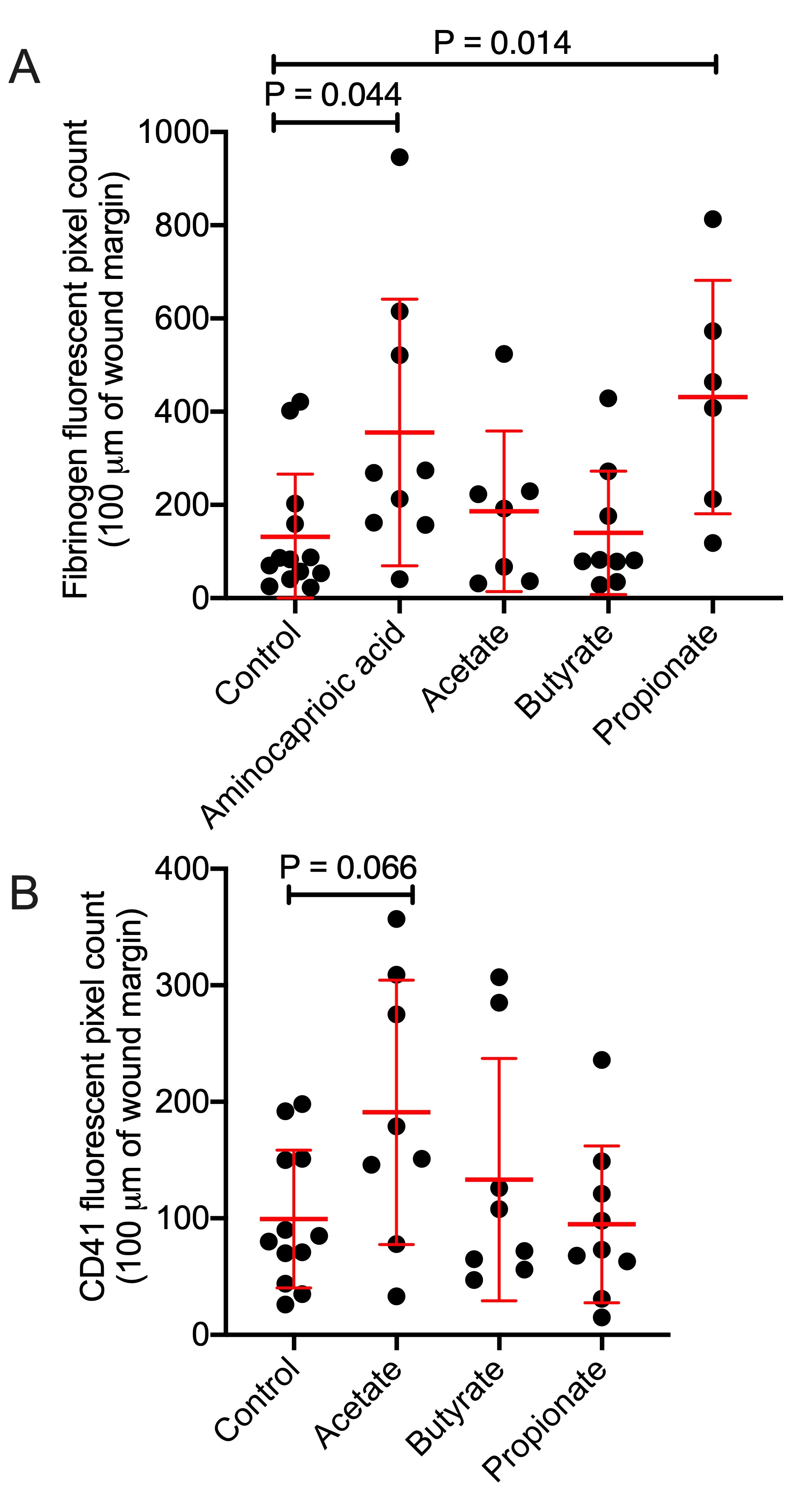

### Supplementary Figure 2

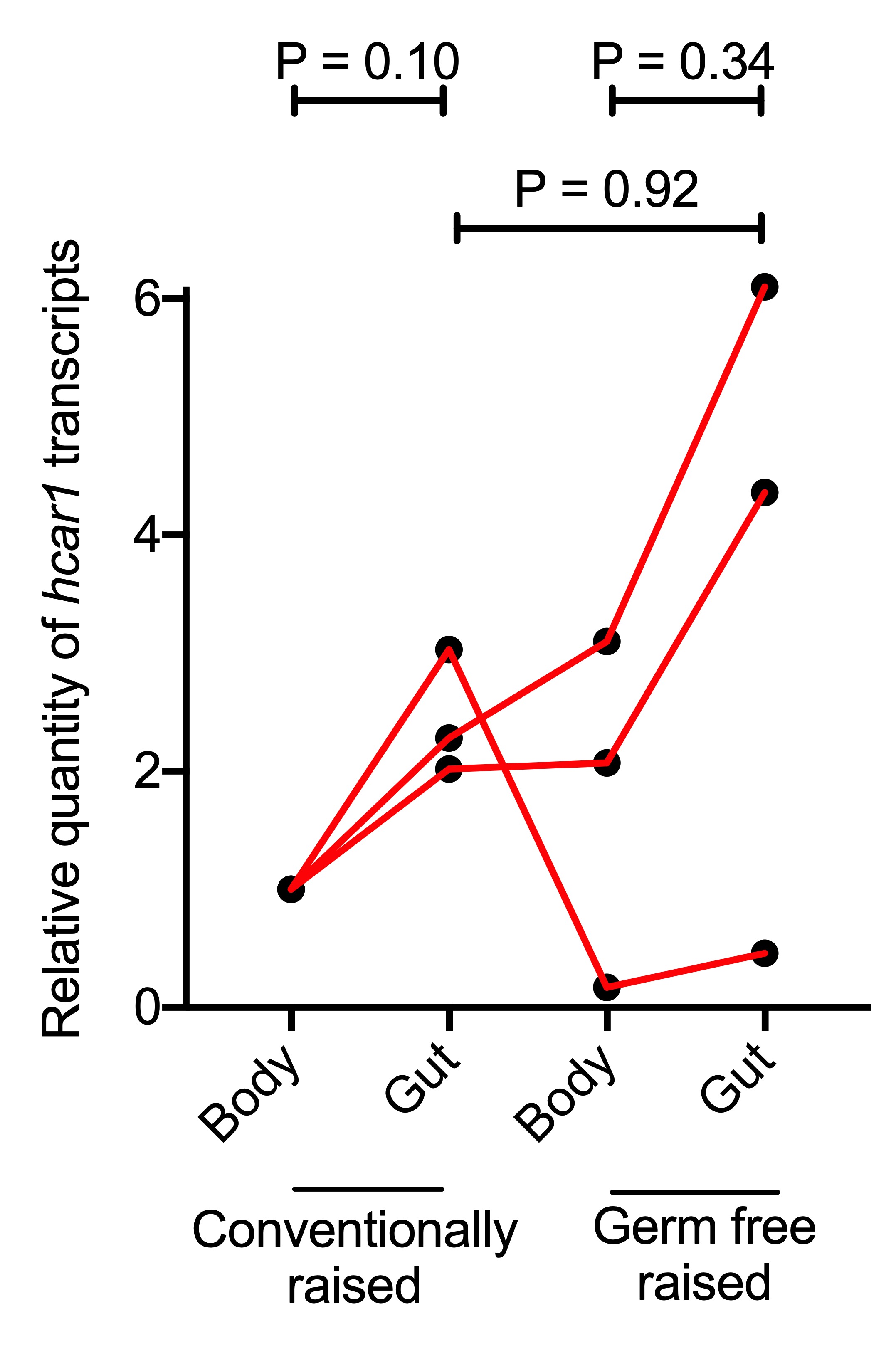
