## Supplementary File for "Conserved anti-inflammatory effects and sensing of butyrate in zebrafish"

```
Zebrafish hcar1a genomic sequence
Hcar1a
>10 dna:chromosome chromosome:GRCz11:10:44449582:44450514:1
ATGAACAACTCCTCATTGTGCTGCACCTATGAAACCCCCCTGCTGGATGCATATCTGCCCCCTGTTCTGTTCAGCGAGTTCGTCCTGGGCCTGATGGGGAACGGTCTGGCCCTCTGCATGTTCTTCTTCCACAGGGACTCCTGGAAGCCCAACTCCATCTATCTGGCTCATTTGGCTCTGGCCAACTCGCTGGTGCTGTTCTGCCTTCCGTTCCGGGCTGACTACTACCGGCGGGGCAAGCACTGGGTCTACGGGGACGCTTTCTGCCGTGTGCTGCTGTTTCTGCTGGCCGCCAATCGAGCCGCCGGTATCTTCTTCCTGACGGCGGTGGCTGTGGACCGATACCTGAAGATCGTGCACCCGCTGAACCGCATCAACCAAATGGGTCTGCGCTACGCCCTGTGGGTGTCTGTGGGCCTGTGGGCGCTCATCATAGCCATGACCGTATATCTGCTCGCCGATAAACACTTCTACTATCTGAACAACCACACTCAATGCGAAAGCTTCAATATTTGCTTTGGGAGCAACTCTCGCTTCACTTGGCATAATGTGTTTTACGTTATTCAATTCTTCGTGCCGACGTTCATCGTCATCTACTGCTCCACATGTATAACCTGGCAGCTGAAGGGTAAAACAATTGACAAGCACGGTAAAATAAGAAGAGCTGTGCGGTTTGTGTTAGCAGTGGCTTTGGTTTTTATCATATGCTTCTTCCCGAGTAACATCTCTCGAATCTCGATGTATGTTCTTAAACTCTGGTATAACGAATGCCAGTATTTTAGTGACGCAAATGATGCATTTAAAACCACAGTCTGCTTTACGTACTTCAACAGCGTGCTCAATCCTGTGGTCTACTACTTCTCCAGCCCTGCCGTCAGCGGATCCTTGAGGAAAATATATATGAGGATTTTAGGACAGAAGATTGAGGAGTGA


Zebrafish hcar1b genomic sequence
>10 dna:chromosome chromosome:GRCz11:10:44457484:44458418:1
ATGAACAACTCCTCCTCGGTGTGCTGCGCTTTCGACGCTCCCATTCTGGATGAGGTGTTACCCCCTGTTCTGTTCAGCGAGTTCGTCCTGGGCCTGATGGGGAACGGTCTGGCCCTCTGCATGTTCTTCTTCCACAGGGACTCCTGGAAGCCCAACTCCATCTATCTGGCTCATCTGGCTCTGGCCGACTCGCTGGTGCTGTTCTGCCTTCCGTTCCGGGCTGACTACTACCGGCGGGGCAAGCACTGGGTCTACGGGGACGCTTTCTGCCGTGTGCTGCTGTTTCTGCTGGCTGCAAATCGAGCCGCCGGTATCTTCTTCCTGACGGCGGTGGCTGTGGACCGATACCTGAAGATCGTGCACCCGCTGAACCCCATCAACCAAATGGGTCTGCGCTACGCCCTGTGGGTGTCTGTGGGCCTGTGGGCGCTCATCATAGCCATGACCGTATATCTGCTCGCCGATAAACACTTCTACTATCGAAATAACCGAACGCAGTGCGAAAGCTTCAATATCTGTTTAGGACACAACGCTCTGTCTGATTGGCACAATTCGTTTTACGTTATTCAATTCTTCGTGCCGACGTTCATCGTCATCTACTGCTCCACATGTATAACCTGGCAGCTGAAGGGTAAAACAATTGACAAGCACGGTAAAATAAGAAGAGCTGTGCGGTTTGTGTTAGCAGTGGCTTTGGTTTTCATCATATGCTTCTTCCCCAGTAATGTCTCCCGTATTGCTGTTTGGGTTCTGAAAAGCTGGCATAATGAATGTCAATATTTTCAGGATGCAAATGTGGCGTTTTATATCACCGTCTGCTTTACGTACTTCAACAGCGTGCTCAATCCTGTGGTCTACTACTTCTCCAGCCCTGCCGTCAGCAGATCCTTGAGGAAAATATATATGAGGATGTCAGGACAGAAGATTGAGGATGA

gRNA target

qFw primer binding site

qRv primer binding site

hcar1a gRNA mismatches to hcar1b sequence
Target 1
A – CAATCGAGCCGCCGGTA
B – AAATCGAGCCGCCGGTA

Target 2 – no clear homology at site

Target 3
A – AGTAACATCTCTCGAATC
B – AGTAATGTCTCCCGTATT

Alignment of hcar1a (top) to hcar1b (bottom)
Matches:864; Mismatches:62; Gaps:16; Unattempted:0

          *         *         *         *         *         *         *         *         *         *
   933<TCACTCCTCAATCTTCTGTCCTAAAATCCTCATATATATTTTCCTCAAGGATCCGCTGACGGCAGGGCTGGAGAAGTAGTAGACCACAGGATTGAGCACG<834
   935<TCA-TCCTCAATCTTCTGTCCTGACATCCTCATATATATTTTCCTCAAGGATCTGCTGACGGCAGGGCTGGAGAAGTAGTAGACCACAGGATTGAGCACG<837


          *         *         *         *         *          *         *         *         *           *
   833<CTGTTGAAGTACGTAAAGCAGACTGTGGTTTTAAATGCATCATTTGCGTCACT-AAAATACTGGCATTCGTTATACCAGAGTTTAAGAA--CATACATCG<737
   836<CTGTTGAAGTACGTAAAGCAGACGGTGATATAAAACGCCACATTTGCATC-CTGAAAATATTGACATTCATTATGCCAGCTTTTCAGAACCCAAACAGC-<739


             *         *         *         *         *         *         *         *         *         *
   736<AGATTCGAGAGATGTTACTCGGGAAGAAGCATATGATAAAAACCAAAGCCACTGCTAACACAAACCGCACAGCTCTTCTTATTTTACCGTGCTTGTCAAT<637
   738<A-ATACGGGAGACATTACTGGGGAAGAAGCATATGATGAAAACCAAAGCCACTGCTAACACAAACCGCACAGCTCTTCTTATTTTACCGTGCTTGTCAAT<640


             *         *         *         *         *         *         *         *         *         *
   636<TGTTTTACCCTTCAGCTGCCAGGTTATACATGTGGAGCAGTAGATGACGATGAACGTCGGCACGAAGAATTGAATAACGTAAAACACATTATGCCAAGTG<537
   639<TGTTTTACCCTTCAGCTGCCAGGTTATACATGTGGAGCAGTAGATGACGATGAACGTCGGCACGAAGAATTGAATAACGTAAAACGAATTGTGCCAA-TC<541


              *          *         *         *         *         *         *         *         *         *
   536<A-AGCGAGAGTTG-CTCCCAAAGCAAATATTGAAGCTTTCGCATTGAGTGTGGTTGTTCAGATAGTAGAAGTGTTTATCGGCGAGCAGATATACGGTCAT<439
   540<AGACAGAGCGTTGTGTCCTAAA-CAGATATTGAAGCTTTCGCACTGCGTTCGGTTATTTCGATAGTAGAAGTGTTTATCGGCGAGCAGATATACGGTCAT<442


               *         *         *         *         *         *         *         *         *         *
   438<GGCTATGATGAGCGCCCACAGGCCCACAGACACCCACAGGGCGTAGCGCAGACCCATTTGGTTGATGCGGTTCAGCGGGTGCACGATCTTCAGGTATCGG<339
   441<GGCTATGATGAGCGCCCACAGGCCCACAGACACCCACAGGGCGTAGCGCAGACCCATTTGGTTGATGGGGTTCAGCGGGTGCACGATCTTCAGGTATCGG<342


               *         *         *         *         *         *         *         *         *         *
   338<TCCACAGCCACCGCCGTCAGGAAGAAGATACCGGCGGCTCGATTGGCGGCCAGCAGAAACAGCAGCACACGGCAGAAAGCGTCCCCGTAGACCCAGTGCT<239
   341<TCCACAGCCACCGCCGTCAGGAAGAAGATACCGGCGGCTCGATTTGCAGCCAGCAGAAACAGCAGCACACGGCAGAAAGCGTCCCCGTAGACCCAGTGCT<242


               *         *         *         *         *         *         *         *         *         *
   238<TGCCCCGCCGGTAGTAGTCAGCCCGGAACGGAAGGCAGAACAGCACCAGCGAGTTGGCCAGAGCCAAATGAGCCAGATAGATGGAGTTGGGCTTCCAGGA<139
   241<TGCCCCGCCGGTAGTAGTCAGCCCGGAACGGAAGGCAGAACAGCACCAGCGAGTCGGCCAGAGCCAGATGAGCCAGATAGATGGAGTTGGGCTTCCAGGA<142


               *         *         *         *         *         *         *         *          *         *
   138<GTCCCTGTGGAAGAAGAACATGCAGAGGGCCAGACCGTTCCCCATCAGGCCCAGGACGAACTCGCTGAACAGAACAGGGGGCAGATA-TGCATCCAGCAG<40
   141<GTCCCTGTGGAAGAAGAACATGCAGAGGGCCAGACCGTTCCCCATCAGGCCCAGGACGAACTCGCTGAACAGAACAGGGGGTA-ACACCTCATCCAGAAT<43


                *         *            *
    39<GGGGGTTTCATAGGTGCAGCACA---ATGAGGAGTTGTTCAT<1
    42<GGGAGCGTCGAAAGCGCAGCACACCGAGGAGGAGTTGTTCAT<1
```

```
Predicted Hcar1a
MNNSSLCCTYETPLLDAYLPPVLFSEFVLGLMGNGLALCMFFFHRDSWKPNSIYLAHLALANSLVLFCLPFRADYYRRGKHWVYGDAFCRVLLFLLAANRAAGIFFLTAVAVDRYLKIVHPLNRINQMGLRYALWVSVGLWALIIAMTVYLLADKHFYYLNNHTQCESFNICFGSNSRFTWHNVFYVIQFFVPTFIVIYCSTCITWQLKGKTIDKHGKIRRAVRFVLAVALVFIICFFPSNISRISMYVLKLWYNECQYFSDANDAFKTTVCFTYFNSVLNPVVYYFSSPAVSGSLRKIYMRILGQKIEE*


Predicted Hcar1b
MNNSSSVCCAFDAPILDEVLPPVLFSEFVLGLMGNGLALCMFFFHRDSWKPNSIYLAHLALADSLVLFCLPFRADYYRRGKHWVYGDAFCRVLLFLLAANRAAGIFFLTAVAVDRYLKIVHPLNPINQMGLRYALWVSVGLWALIIAMTVYLLADKHFYYRNNRTQCESFNICLGHNALSDWHNSFYVIQFFVPTFIVIYCSTCITWQLKGKTIDKHGKIRRAVRFVLAVALVFIICFFPSNVSRIAVWVLKSWHNECQYFQDANVAFYITVCFTYFNSVLNPVVYYFSSPAVSRSLRKIYMRMSGQKIED


Alignment Hcar1a and Hcar1b amino acid sequence
Identities:275/309(89%), Positives:289/309(93%), Gaps:0/309(0%)

Hcar1a 2    NNSSLCCTYETPLLDAYLPPVLFSEFVLGLMGNGLALCMFFFHRDSWKPNSIYLAHLALA  61
            N+SS+CC ++ P+LD  LPPVLFSEFVLGLMGNGLALCMFFFHRDSWKPNSIYLAHLALA
Hcar1b 3    NSSSVCCAFDAPILDEVLPPVLFSEFVLGLMGNGLALCMFFFHRDSWKPNSIYLAHLALA  62

Hcar1a 62   NSLVLFCLPFRADYYRRGKHWVYGDAFCRVLLFLLAANRAAGIFFLTAVAVDRYLKIVHP  121
            +SLVLFCLPFRADYYRRGKHWVYGDAFCRVLLFLLAANRAAGIFFLTAVAVDRYLKIVHP
Hcar1b 63   DSLVLFCLPFRADYYRRGKHWVYGDAFCRVLLFLLAANRAAGIFFLTAVAVDRYLKIVHP  122

Hcar1a 122  LNRINQMGLRYALWVSVGLWALIIAMTVYLLADKHFYYLNNHTQCESFNICFGSNSRFTW  181
            LN INQMGLRYALWVSVGLWALIIAMTVYLLADKHFYY NN TQCESFNIC G N+   W
Hcar1b 123  LNPINQMGLRYALWVSVGLWALIIAMTVYLLADKHFYYRNNRTQCESFNICLGHNALSDW  182

Hcar1a 182  HNVFYVIQFFVPTFIVIYCSTCITWQLKGKTIDKHGKIRRAVRFVLAVALVFIICFFPSN  241
            HN FYVIQFFVPTFIVIYCSTCITWQLKGKTIDKHGKIRRAVRFVLAVALVFIICFFPSN
Hcar1b 183  HNSFYVIQFFVPTFIVIYCSTCITWQLKGKTIDKHGKIRRAVRFVLAVALVFIICFFPSN  242

Hcar1a 242  ISRISMYVLKLWYNECQYFSDANDAFKTTVCFTYFNSVLNPVVYYFSSPAVSGSLRKIYM  301
            +SRI+++VLK W+NECQYF DAN AF  TVCFTYFNSVLNPVVYYFSSPAVS SLRKIYM
Hcar1b 243  VSRIAVWVLKSWHNECQYFQDANVAFYITVCFTYFNSVLNPVVYYFSSPAVSRSLRKIYM  302

Hcar1a 302  RILGQKIEE  310
            R+ GQKIE+
Hcar1b 303  RMSGQKIED  311
```
